# Aberrant FGF signaling promotes granule neuron precursor expansion in SHH subgroup infantile medulloblastoma

**DOI:** 10.1101/2021.06.23.449636

**Authors:** Odessa R. Yabut, Hector Gomez, Jessica Arela, Jesse Garcia Castillo, Thomas Ngo, Samuel J. Pleasure

**Affiliations:** Department of Neurology, Weill Institute for Neuroscience, University of California San Francisco, San Francisco, California; Department of Psychiatry, Weill Institute for Neuroscience, University of California San Francisco, San Francisco, California, USA; Programs in Neuroscience and Developmental Biology, Eli and Edythe Broad Center of Regeneration Medicine and Stem Cell Research, University of California San Francisco, California, USA

## Abstract

Mutations in Sonic Hedgehog (SHH) signaling pathway genes, e.g., *Suppressor of Fused* (SUFU), drive granule neuron precursors (GNP) to form medulloblastomas (MB^SHH^). However, how different molecular lesions in the Shh pathway drive transformation is frequently unclear, and *SUFU* mutations in the cerebellum seem distinct. In this study, we show that fibroblast growth factor 5 (FGF5) signaling is integral for many infantile MB^SHH^ cases and that *FGF5* expression is uniquely upregulated in infantile MB^SHH^ tumors. Similarly, mice lacking SUFU (Sufu-cKO) ectopically express *Fgf5* specifically along the secondary fissure where GNPs harbor preneoplastic lesions and show that FGFR signaling is also ectopically activated in this region. Treatment with an FGFR antagonist rescues the severe GNP hyperplasia and restores cerebellar architecture. Thus, direct inhibition of FGF signaling may be a promising and novel therapeutic candidate for infantile MB^SHH^.

## Introduction

Medulloblastoma (MB) is the most common malignant brain tumor in children, with half of cases diagnosed before the age of 5 (Ward et al., 2014; Ostrom et al., 2016). Mutations in *Suppressor of Fused* (*SUFU)* comprise approximately 30% of tumors in infants with Sonic Hedgehog-driven MB (MB^SHH^). Infantile MB^SHH^, including those associated with *SUFU* mutations (MB^SHH-SUFU^), has a worse prognosis and higher rates of local recurrence than other MB^SHH^ subtypes (Kool et al., 2014; Schwalbe et al., 2017; Guerrini-Rousseau et al., 2018). Unfortunately, available SHH-targeted treatments for MB^SHH^ act specifically on proteins upstream of SUFU, and are therefore ineffective for MB^SHH-SUFU^ patients (Kool et al., 2014). The poor prognosis, early occurrence, and lack of targeted therapy for MB^SHH-SUFU^ patients make a detailed understanding of the drivers of oncogenesis in this group of great importance.

SUFU acts as an intracellular modulator of SHH signaling (Matise and Wang, 2011). Briefly, the SHH signaling pathway is initiated after the binding of extracellular Shh ligands to the transmembrane receptor Patched 1 (PTCH1). This relieves PTCH1 inhibition of the transmembrane protein, Smoothened (SMO), and enables the initiation of a cascade of intracellular events promoting the activator function of the transcription factors, GLI1, GLI2, or GLI3. SUFU modulates SHH signaling by ensuring the stability of GLI transcription factors or by promoting the formation of the repressor forms of GLI2 (GLI2R) or GLI3 (GLI3R) (Chen et al., 2009; Wang et al., 2010; Lin et al., 2014). Thus, depending on the developmental context, loss of SUFU can lead to activation or repression of SHH signaling. In the developing cerebellum, SUFU dysfunction is associated with the abnormal development of granule neuron precursors (GNP), which account for MB^SHH^ (Kim et al., 2011, 2018; Vanner et al., 2014; Kong et al., 2019; Vladoiu et al., 2019; Yin et al., 2019; Jiwani et al., 2020). GNPs populate the external granule layer (EGL) along the cerebellar perimeter, where local SHH signals trigger GNP proliferation and differentiation at neonatal stages. However, other mitogenic pathways can also influence GNP behavior (Leto et al., 2016), yet little is known about how SUFU interacts with these pathways. Understanding how *Sufu* loss-of-function (LOF) affects the activity of concurrent local signaling pathways in granule neuron development may be key to developing potent targets for anti-tumor therapy.

In this study, we examined the regulation of FGF signaling in MB^SHH^ and identified upregulation of *FGF5* expression in tumors of infantile MB^SHH^ patients. Similarly, we show ectopic expression of *Fgf5* in the neonatal cerebellum of mice lacking SUFU, which correlates with the activation of FGF signaling in surrounding EGL-localized cells where GNPs accumulate. Strikingly, acute pharmacological inhibition of FGF signaling results in near-complete rescue of these defects, including restoration of cerebellar histo-organization. Thus, our findings identify *FGF5* as a potential biomarker for a subset of patients with infantile MB^SHH^ who may be responsive to FGFR-targeting therapies.

## Results

### *FGF5* is specifically upregulated in SHH-driven infantile MB

We previously reported that *Sufu* LOF in neocortical progenitors results in FGF signaling activation to influence the specification and differentiation of neocortical excitatory neurons (Yabut et al., 2020). Thus, we sought to determine if key FGF signaling pathway genes are differentially expressed in MB patient tumors. We performed a comparative analysis of the expression dataset from 763 MB patient samples comprised of tumors resected from molecularly distinct MB subgroups: Wingless (Wnt subgroup; MB^WNT^), MB^SHH^, Group 3 (MB^Group3^), and Group 4 (MB^Group4^) (Cavalli et al., 2017). Strikingly, our analyses show that *FGF5* expression is higher in tumors, specifically from MB^SHH^ patients, compared to other MB subgroups, with approximately 25% of MB^SHH^ tumors exhibiting a two-fold increase (**Fig. 1A-1B**). We also find that *FGF5* is uniquely upregulated in MB^SHH^ tumors from patients within the 0-3 years old age group, but not patients within the same age group in other MB subtypes (**Fig. 1C**). Further examination across all MB^SHH^ tumors stratified which subgroups express the highest levels of *FGF5* expression. Infantile tumors, largely belonging to the SHHb and SHHg subgroups (Cavalli et al., 2017), exhibit higher *FGF5* levels compared to tumors from children (SHHa) or adults (SHHd) (**Fig. 1D**). By all measures, the proportion of SHHb and SHHg tumors with relatively high levels of *FGF5* is significantly increased (∼30%) compared to other MB^SHH^ subgroups (**Fig. 1E**). Taken together, these findings strongly suggest that FGF signaling is specifically disrupted in infantile-onset MB^SHH^.

**Figure 1:**
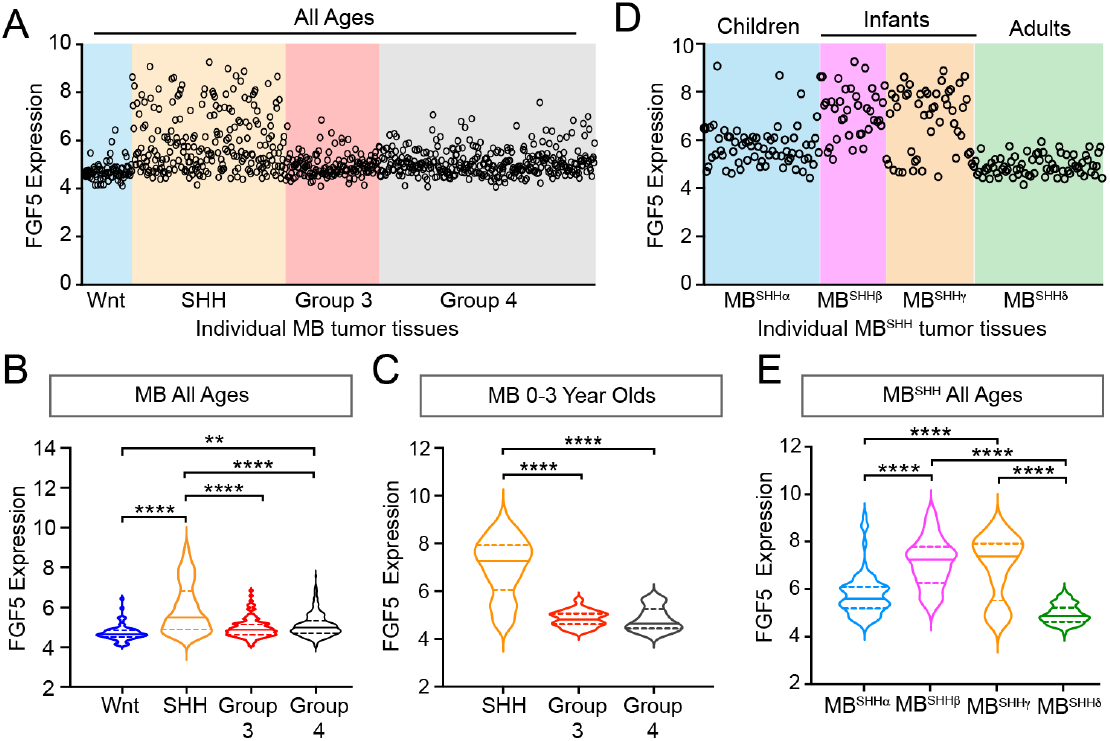
Upregulated *FGF5* expression in MB^SHH^ tumors from infant patients. **(A)** Levels of FGF5 expression in human MB tumors of all ages from GEO expression dataset #GSE85217(Cavalli et al., 2017). **(B, C)** Statistical analysis of *FGF5* expression levels associated with MB tumor subtypes from patients across all ages (B) and 0-3 year old MB patients (C). **p<0.01, ****p<0.0001. **(D, E)** Graph represents *FGF5* expression levels in human MB^SHH^ tumors of all ages from GEO expression dataset #GSE85217 (D) and corresponding plots (E) showing statistically higher *FGF5* expression in tumors from infants with MB^SHH^ compared to tumors from children or adults with MB^SHH^. ****p<0.0001.

### Region-specific expansion of GNPs in the P0 Sufu-cKO cerebellum coincides with increased *Fgf5* expression

FGF signaling has been implicated in cerebellar development, particularly in granule neuron development (Yaguchi et al., 2009; Yu et al., 2011), leading us to wonder if and how aberrant FGF signaling may be contributing to the oncogenicity of GNPs. Since mutations in *SUFU* drive infantile MB^SHH^, we generated the mutant mice (*hGFAP-Cre;Sufu*^*fl/fl*^, hereto referred to as Sufu-cKO), in which *Sufu* is conditionally deleted in GNPs (Zhuo et al., 2001) to examine FGF activity. Sufu-cKO mice exhibit profound defects in cerebellar development. At P0, a time point at which GNP proliferation and differentiation is ongoing, there is a visible increase in measured cerebellar size and expansion of Pax6-positive (Pax6+) GNPs in the EGL of Sufu-cKO cerebellum compared to controls (**Fig. 2A-C**). Notably, the expansion of Pax6+ GNPs specifically localizes along the secondary fissure (EGL Region B, arrow in **Fig. 2A**) compared to other EGL areas (EGL Regions A and C) in the P0 Sufu-cKO cerebellum (**Fig. 2D**). We then proceeded to examine whether these defects correlated with abnormal FGF5 expression. We collected P0 littermates for these analyses and found that the foliation defects are particularly severe in the cerebelli of mice from this P0 Sufu-cKO litter compared to littermate controls (as determined by DAPI labeling in **Fig. 2E**). *In situ* hybridization (ISH) using *Fgf5*-specific riboprobes show high expression of *Fgf5* (Fgf5^high^) immediately adjacent to the presumptive secondary fissure in the P0 control cerebellum (**Fig. 2E**). Strikingly, in the P0 Sufu-cKO cerebellum, there is an expansion of Fgf5^high^ expression regions (**Fig. 2E**), coinciding with areas near the secondary fissure where GNP expansion is most severe (**Fig. 2A, 2D**). Further, while *Fgf5*-expressing (FGF5+) cells are largely excluded from the EGL of the control cerebellum, a substantial number of *Fgf5*+ cells are ectopically localized in the EGL Region B of the Sufu-cKO cerebellum (**Fig. 2E, 2F**). *Fgf5* mRNA molecules are visibly higher in *Fgf5*-expressing cells in the Sufu-cKO EGL, while *Fgf5* expression is largely absent in the deeper EGL regions (outlined cells within boxed regions in **Fig. 2F**). These findings implicate FGF5 as a potential instigator of region-specific defects in GNP differentiation present in the P0 Sufu-cKO cerebellum.

**Figure 2:**
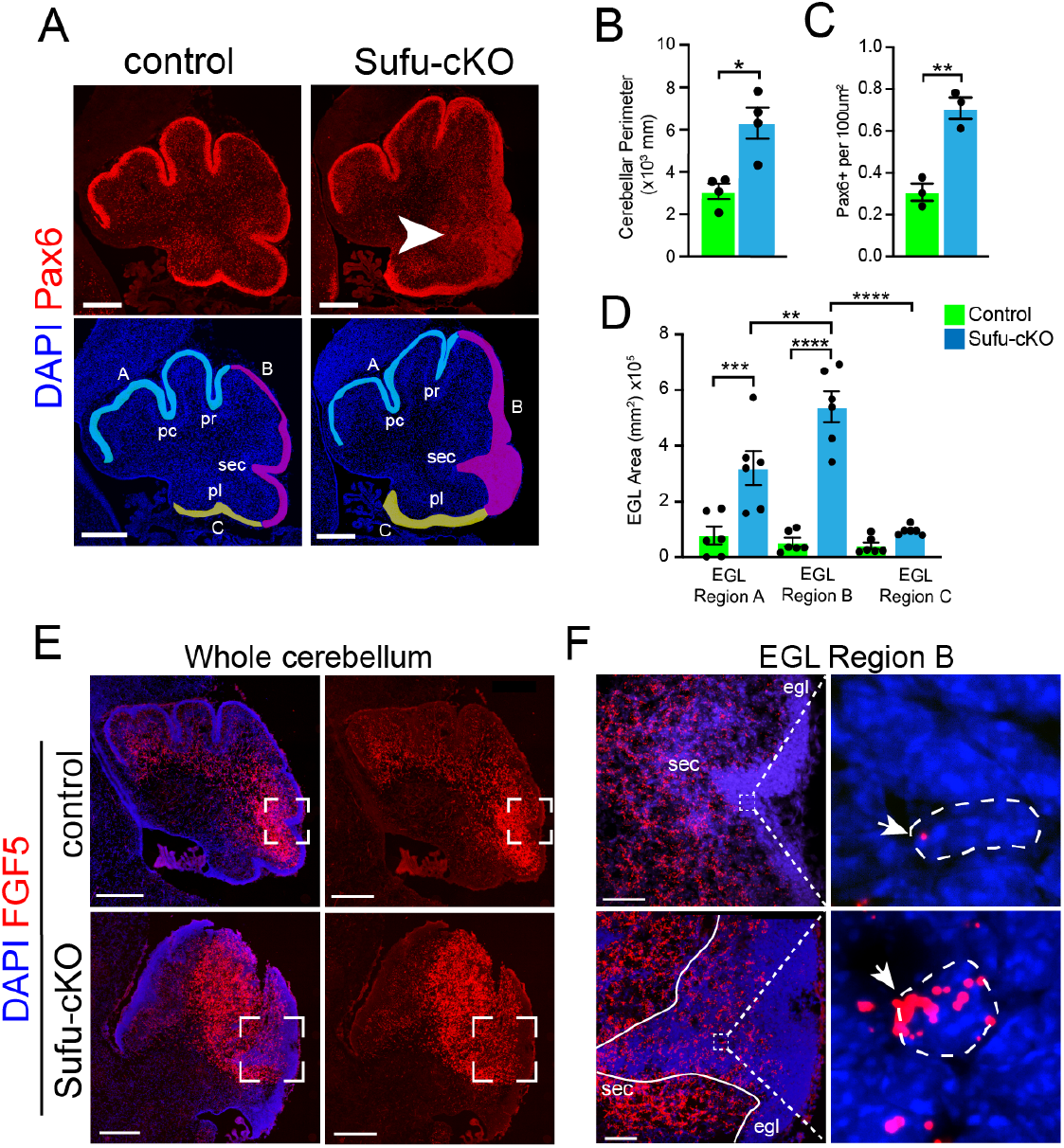
Increased FGF5 expression coincides with region-specific expansion of GNPs in the P0 Sufu-cKO cerebellum. **(A)** Pax6 (red) and DAPI (blue) immunofluorescence staining of the P0 Sufu-cKO and control cerebelli. Arrow points to severely expanded EGL region B in the P0 Sufu-cKO cerebellum. EGL regions are designated in DAPI-labeled sections as A (light blue), B (magenta), and C (yellow). Each region encompasses specific fissures: the preculminate (pc) and primary (pr) fissures for Region A, the secondary (sec) fissure for Region B, and the posterolateral (pl) fissure for Region Scale bars: Scale bars = 250 mm. **(B-D)** Quantification and comparison of the cerebellar perimeter (B), total area occupied by densely packed Pax6+ cells (C), and size of specific EGL regions (D) between P0 Sufu-cKO and control cerebelli. **(E)** Fluorescent *in situ* hybridization using RNAScope probes against *Fgf5* mRNA (red) shows the expansion of *Fgf5* expression in the P0 Sufu-cKO cerebellum compared to controls. Sections are counterstained with DAPI to distinguish structures. Boxed areas are magnified in F. Scale bars = 500 mm. **(F)** *Fgf5* is ectopically expressed in cells within the EGL of the P0 Sufu-cKO cerebellum. Boxed areas within the EGL show DAPI-labeled cells expressing visibly high levels of *Fgf5*, identified as punctate labeling (arrowheads), in the EGL of P0 Sufu-cKO cerebellum compared to controls. Scale bars = 50 mm. \*\**p*<0.05, **p<0.01, ***p<0.001, ****p<0.0001.

### FGF signaling drives GNP proliferation in the P0 Sufu-cKO cerebellum

FGF5 is a ligand for fibroblast growth factor receptors 1 (FGFR1) and 2 (FGFR2), both of which are expressed in the developing cerebellum, particularly in IGL regions where *Fgf5*-expressing cells localize (Clements et al., 1993; Ornitz et al., 1996). The binding of FGF5 to these receptors triggers the activation of multiple intracellular signaling pathways, including the mitogen-activated protein kinase (MAPK) pathway, to control cellular activities driving NSC progression (**Fig. 3A**) (Ornitz and Itoh, 2015). Immunostaining with antibodies to detect MAPK pathway activation reveals increased MAPK activity in the EGL of the P0 Sufu-cKO cerebellum. Regional distribution of cells labeled with phospho-Erk1/2 (pErk1/2+), a marker for activated MAPK signaling, shows that the abnormally expanded EGL of the secondary fissure in the P0 Sufu-cKO cerebellum increase in these cells (**Fig. 3B**). We quantified the number of pERK1/2+ cells in proliferative regions defined by dense Ki-67-labeled cells and found a significant increase in pERK1/2+ cells compared to controls (**Fig. 3C**). Many pERK1/2+ cells are also proliferative as indicated by Ki-67 labeling (boxed areas, **Fig. 3B**) and the numbers of these dual labeled cells were significantly higher in the Sufu-cKO cerebellum compared to controls (**Fig. 3D**). These findings indicate that the increase in *Fgf5* expression correlates with the activation of FGF signaling in GNPs and demonstrates a likely role in regulating the abnormal proliferation and pre-neoplastic lesion in mutant mice.

**Figure 3:**
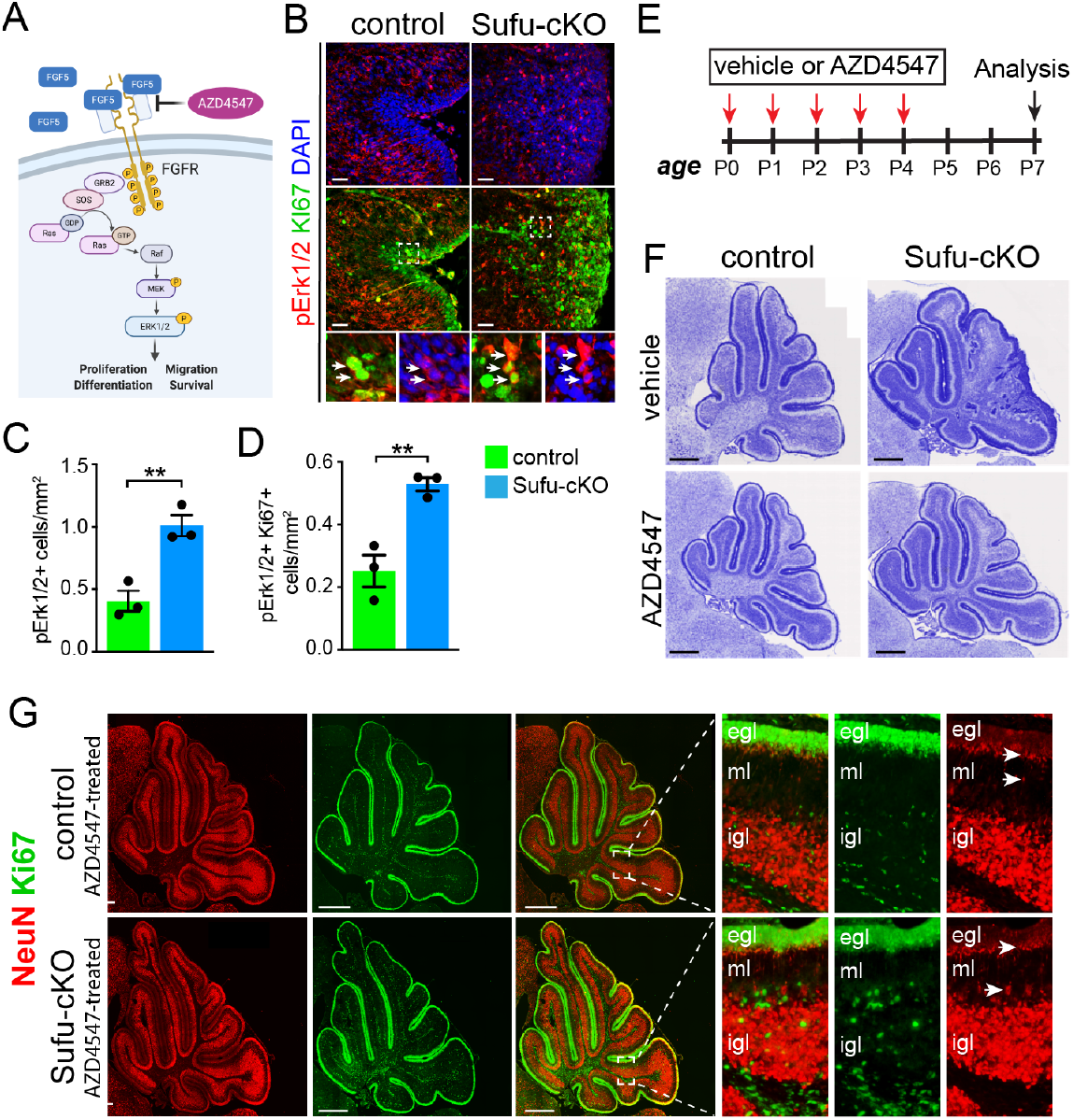
Ectopic activation of FGF signaling in the EGL of P0 Sufu-cKO cerebellum. **(A)** Schematic diagram showing the activation of FGF signaling activity upon binding of FGF5 to extracellular domains of FGFR via the MAPK signal transduction pathway. **(B)** Double-immunofluorescence staining with Ki-67 (green) and phospho-Erk1/2 (pErk1/2; red), a marker of activated MAPK signaling in the P0 Sufu-cKO and control cerebelli. Boxed regions show pErk1/2+ and Ki-67+ cells (arrowheads) in the control and Sufu-cKO EGL. Scale bars = 50 μm. **(C, D)** Quantification of pErk1/2+ cells (C) and double-labeled pErk1/2+ and Ki-67+ cells (D) in the P0 Sufu-cKO and control EGL Region B. **p<0.01. **(E)** Experimental design of rescue studies performed by intraventricular administration of FGFR1-3 pharmacological inhibitor, AZD4547, or vehicle controls. **(F)** Nissl staining of the P7 control and Sufu-cKO treated with either AZD4547 or vehicle, 2 days after treatment. Scale bars = 500 μm. **(G)** NeuN and Ki-67 double immunofluorescence staining of the P7 control and Sufu-cKO treated with AZD4547. Boxed regions show localization and organization of NeuN+ and Ki-67+ cells in distinct cerebellar layers. Arrows point to areas of the EGL and IGL where NeuN+ cells are beginning to be expressed.

Given the role of SUFU in regulating GLI transcription factors to modulate SHH signaling activity (Kim et al., 2018; Yin et al., 2019), we examined whether the activation of FGF signaling occurred ^C.^ concurrently with SHH signaling activation. Surprisingly, GLI protein levels in the P0 control and Sufu-cKO cerebellum show a marked reduction of GLI1, GLI2, GLI3, and PTCH1 levels (**Supplementary Fig. 1A**); this is inconsistent with elevated SHH signaling we anticipated in the absence of SUFU. To directly examine this in specific EGL regions, we compared the cerebellum of P0 control and Sufu-cKO mice carrying the Shh reporter transgene *Gli1-LacZ* (Bai et al., 2002). In these mice, Shh signaling activity is absent or very low throughout the entire P0 Sufu-cKO cerebellum but is highly active in a region-specific manner in controls (**Supplementary Fig. 1B**). Furthermore, while some LacZ+ cells are detectable in EGL regions A and C and adjacent ML and IGL, LacZ+ cells are completely absent in EGL region B and adjacent areas of the P0 Sufu-cKO cerebellum (**Supplementary Fig. 1B-1C**). These findings indicate that the accumulation of GNPs does not rely on active Shh signaling, particularly in Region B, where there is a severe expansion of GNPs in the absence of SUFU.

### Blockade of FGF signaling dramatically rescues the Sufu-cKO phenotype

To determine if activated FGF signaling drives GNP defects in the Sufu-cKO cerebellum, we pharmacologically inhibited FGF signaling using the competitive FGFR1-3 antagonist AZD4547 (Gudernova et al., 2016). For this experiment, 1 μl of AZD4547 (5mg/ml) was delivered via intraventricular (IV) injection for 5 consecutive days beginning at P0, and the cerebellum was analyzed 3 days after treatment at P7 (**Fig. 3E**). Strikingly, AZD4547 treatment results in near complete rescue of the GNP phenotype by P7 in the Sufu-cKO cerebellum, displaying a cerebellar morphology indistinguishable from controls with normal foliation and cellular organization (**Fig. 3F**). Indeed, in the cerebellum of AZD4547 -treated P7 Sufu-cKO mice, proliferating Ki-67+ cells largely exclusively localize in the EGL while NeuN+ cells densely pack the IGL and not the EGL (**Fig. 3G**). Notably, NeuN expression appears in cells localized at the border of the EGL and ML, where Ki-67+ cells are absent, indicating that post-mitotic cells successfully began differentiation as observed in controls (boxed regions, **Fig. 3G**). Thus, our findings confirm that inhibition of FGF signaling in proliferating GNPs of the Sufu-cKO cerebellum ensure normal progression of GNP differentiation.

### *Fgf5* expression is increased in the developing cerebellum of Sufu;p53-dKO mice

Our findings indicate a critical role for FGF signaling in driving GNP hyperplasia, making GNPs vulnerable to neoplastic lesions, resulting in tumorigenesis when SUFU is absent in the developing cerebellum. Indeed, we find that Pax6+ GNPs within the neonatal Sufu-cKO EGL display an increase in double-strand breaks, especially in Region B (DSBs; **Fig. 4A-4B**), as detected by immunostaining for phosphorylated H2AX (γH2AX), an early marker for DSBs (Mah et al., 2010). Nevertheless, as previously reported, tumors do not readily form in the Sufu-cKO cerebellum (Yin et al., 2019), indicating either timely repair of DSBs or the induction of apoptosis in GNPs with significant genomic instability. Indeed, double-labeling with cleaved Caspase 3 (CC3) and γH2AX shows a significantly higher number of double-labeled cells in EGL Regions A and B (**Supplementary Fig. 2A-2B**). Among the downstream targets of DSB repair pathways is p53, which, when activated, mediates cell death to suppress tumor formation (Gao et al., 2000). In the P0 Sufu-cKO cerebellum, p53 protein is present, albeit significantly reduced (**Fig. 4C**). The reduction in p53 may be driving an increase in DSBs, yet it is still sufficient to induce apoptotic pathways. Supporting this, conditional ablation of both *p53* and *Sufu* in GNPs (*hGFAP-Cre;Sufu*^*fl/fl*^*;p53*^*fl/fl*^ or Sufu;p53-dKO) results in the formation of massive tumors in the cerebellum within 2 months after birth (**Fig. 4F**), indicating the failure to activate critical apoptotic pathways. However, tumors do not form in mice lacking *p53* (Marino et al., 2000) or *Sufu* alone (**Fig. 4F**). These findings suggest that in the absence of SUFU, the failure of GNPs to transition into fully differentiated granule neurons compromises genomic stability and renders GNPs extremely vulnerable to tumor formation with a second molecular hit.

**Figure 4:**
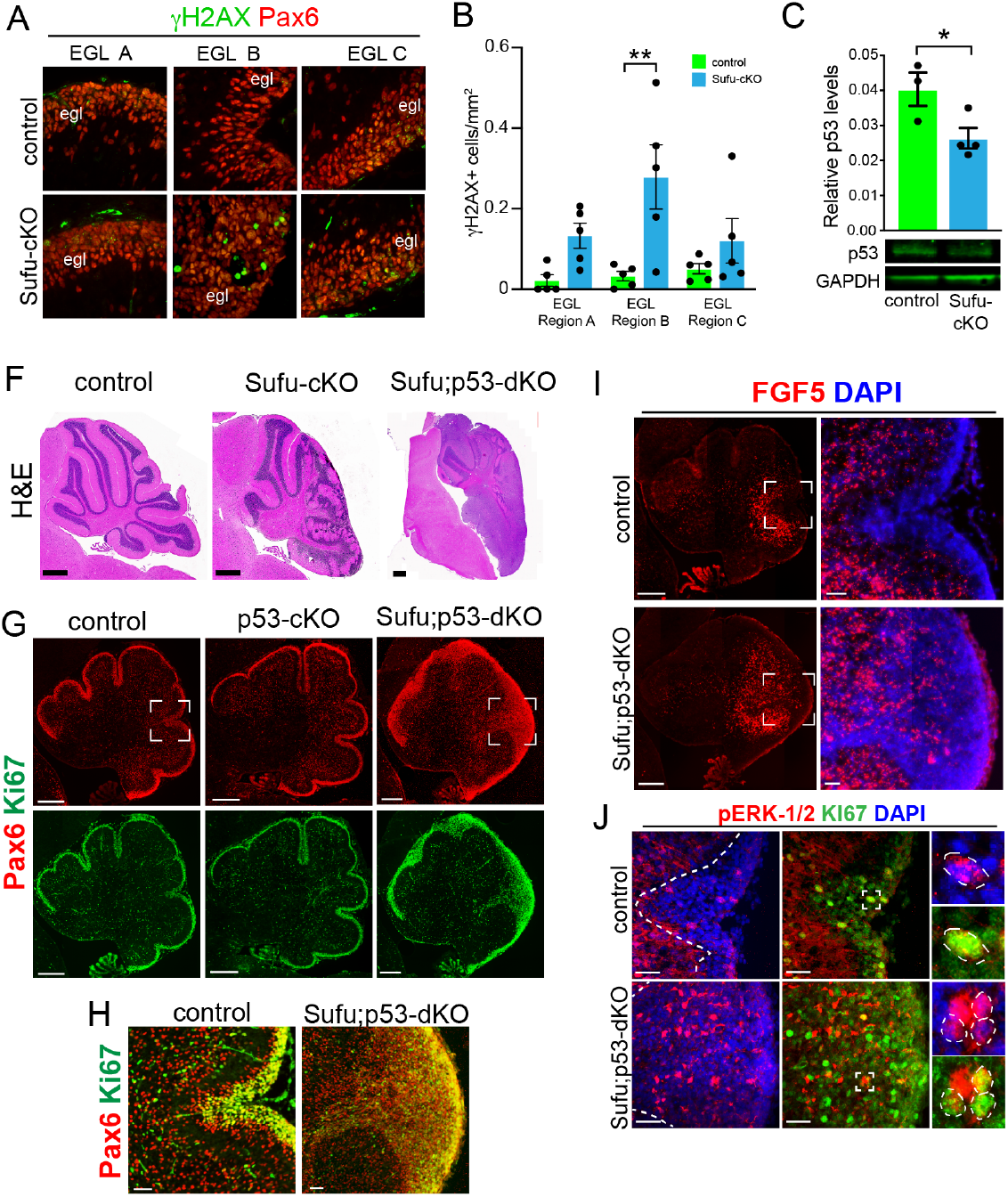
Evidence of pre-neoplastic lesions and high rates of cell death in Sufu-cKO granule neuron precursors. **(A)** Double-immunofluorescence staining with Pax6 (red) and (γH2AX (green), a marker for double-strand DNA breaks in specific EGL regions of the P0 Sufu-cKO and control cerebella. **(B)** Quantification of (γH2AX + cells in each cerebellar region of P0 control and Sufu-cKO mice. **p<0.01. **(C)** Western blot analysis of p53 protein levels in P0 control and Sufu-cKO cerebellar protein lysates. \**p*<0.05. **(F)** Hematoxylin and Eosin (H&E) staining of P60 control, Sufu-cKO, Sufu;p53-dKO cerebella. Scale bars = 500 μm. **(G, H)** Double-immunofluorescence staining against Pax6 (red) and Ki-67 (green) in the P0 control, p53-cKO, and Sufu;p53-dKO cerebellum (G). Boxed regions in G are magnified in H, demonstrating the expansion of the EGL in the P0 Sufu;p53-dKO cerebellum compared to littermate controls. Scale bars = 200 μm (A) and 50 μm (B). **(I)** Fluorescent *in situ* hybridization using RNAScope probes against *Fgf5* mRNA (red) and DAPI labeling in the P0 Sufu;p53-dKO and control cerebellum. Boxed areas are enlarged to show ectopic localization of *Fgf5*+ cells in the EGL of the Sufu-p53-dKO cerebellum, unlike in controls. Scale bars = 200 μm and 50 μm (boxed area). **(J)** Double-immunofluorescence staining with Ki-67 (green) and phospho-Erk1/2 (pErk1/2; red) in the P0 Sufu;p53-dKO and control cerebelli. Boxed regions show cells double-labeled with pErk1/2+ and Ki-67+ cells in the control and Sufu;p53-dKO EGL Region B. Scale bars = 25 μm.

We sought to confirm that upregulated FGF signaling also occurs in the tumor-prone Sufu;p53-dKO. As expected, the neonatal cerebellum of Sufu-cKO displays EGL expansion due to excess proliferative (Ki-67+) and Pax6+ cells in the P0 Sufu;p53-dKO cerebellum but not in p53-cKO and control cerebelli (**Fig. 4G**). Further expansion of Pax6+ GNPs in the P0 Sufu;p53-dKO is also most severe along the secondary fissure (EGL Region B) compared to other EGL areas (EGL Regions A and C) of the P0 Sufu;p53-dkO cerebellum (**Fig. 4G-4H**). As with our observations in the P0 Sufu-cKO cerebellum, ISH for *Fgf5* in the P0 Sufu;p53-dKO cerebellum shows ectopic *Fgf5* expression. Particularly, *Fgf5*+ cells are expanded anteriorly and detected specifically around the secondary fissure of the P0 Sufu;p53-dKO cerebellum (**Fig. 4I**). There is also ectopic MAPK signaling activity in the P0 Sufu;p53-dKO cerebellum, with significantly higher numbers of pErk1/2+ cells within the EGL, many of which are proliferative as marked by co-labeling with Ki-67, within the expanded EGL of the secondary fissure (**Fig. 4J**). These findings indicate that, as in the P0 Sufu-cKO cerebellum, ectopic *Fgf5* expression triggers FGF signaling in GNPs in the P0 Sufu;p53-dKO cerebellum and may facilitate oncogenic transformation and tumor growth of GNPs.

## Discussion

Our studies identify a mechanism by which the combinatorial effects of oncogenic *SUFU* mutations and other concurrent developmental signaling pathways make GNPs vulnerable to oncogenic transformation leading to infantile MB^SHH^. Using mice lacking *Sufu* in GNPs, we find that ectopic *Fgf5* expression correlates with an increase in FGF signaling, particularly in areas where proliferating GNPs reside, resulting in GNP hyperplasia, preneoplastic lesions, and patterning defects. Inhibition of Fgf signaling through pharmacological blockade of FGFR1-3 prevents hyperplasia and associated cerebellar architectural abnormalities. Strongly supporting a role for FGF5, we also find elevated levels of *Fgf5* gene expression specifically in infantile MB^SHH^ patients, but not in other MB subgroups. These findings indicate that FGF-targeting compounds may be a promising therapeutic option for infantile MB^SHH^ patients, with elevated levels of FGF5 in tumor tissues acting as a potential biomarker.

Expansion of Pax6+ GNPs in the newborn Sufu-cKO cerebellum (**Fig. 2**) occurs in the posterior/ventral regions of the cerebellar hemispheres where infantile MB tumors typically arise (Tan et al., 2018). Interestingly, these subregions have low levels of GLI1 reporter activity, PTCH1 expression, and SHH ligand expression (Corrales et al., 2004). Spatially distinct regulation of granule neuron development by SUFU may, therefore, rely on non-canonical SHH signaling beyond SMO or through yet undefined downstream interactions, resulting in control of FGF signaling activity. Supporting this, we find ectopic expression of *Fgf5* and deregulation of FGF signaling as marked by the excessive intracellular activation of MAPK signaling in the P0 Sufu-cKO cerebellum. Similarly, in previous studies, ectopic expression of FGF ligands, *Fgf8* and *Fgf15*, are associated with degradation of GLI3R because of *Sufu* deletion, resulting in regional patterning and precursor specification and differentiation defects (Kim et al., 2011, 2018; Jiwani et al., 2020; Yabut et al., 2020). Taken together, these findings indicate that SUFU acts at the intersection of SHH and FGF signaling to modulate GNP behavior (as summarized in our working model in **Fig. 5**). We postulate that SUFU exerts this role via stabilization of GLI transcription factors since we observed a decrease in GLI1, GLI2, and GLI3 protein level in the newborn Sufu-cKO cerebellum. Supporting this, Yin et al. (2019) found that reducing GLI2 levels rescued GNP hyperplasia and patterning defects in the Sufu-cKO neonatal cerebellum. Further studies are required to elucidate the involvement of GLI-dependent mechanisms in controlling *Fgf5* gene expression in the neonatal cerebellum.

**Figure 5:**
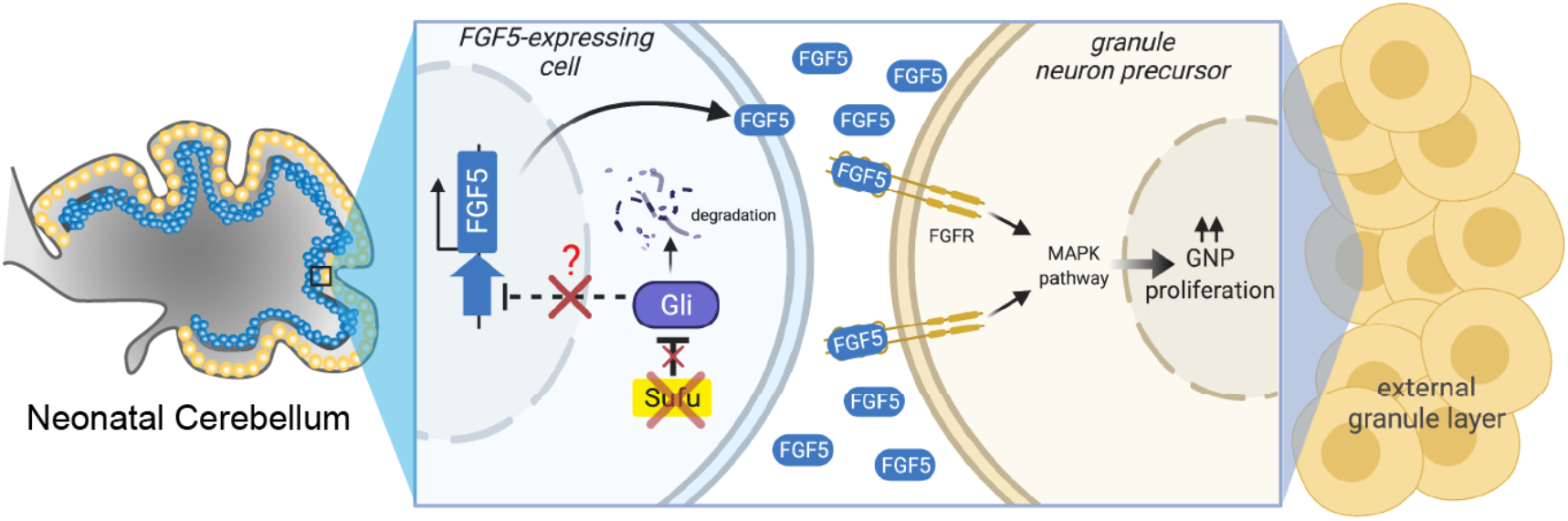
Loss of *Sufu* function drives excess proliferation of granule neuron precursors via FGF signaling activation. The schematic diagram models how *Sufu* LOF facilitates the expansion of GNPs (yellow cells) in the EGL at the early stages of cerebellar development. We hypothesize that *Fgf5*-expressing cells (blue cells, yet to be identified) send FGF5 signal to GNP to proliferate and that ectopic expression of *Fgf5* in the absence of SUFU is responsible for the uncontrolled expansion of EGL localized GNPs.

In the wildtype cerebellum, *Fgf5*-expressing cells localize within the IGL surrounding the secondary fissure at P0 (**Fig. 4B**) (Ozawa et al., 1996; Hattori et al., 1997; Yaguchi et al., 2009) where GLI1 reporter activity (**Fig. 4A**), and SHH and PTCH1 expression are lowest (Corrales et al., 2004). By P4, *Fgf5* expression is detected in cells within the IGL throughout the cerebellum (Yaguchi et al., 2009). However, by P14, a time point at which most granule neurons have differentiated, *Fgf5* is no longer detected in the IGL or other cerebellar regions per Allen Developing Mouse Brain Atlas (Allen Institute for Brain Science, 2004). The strict spatiotemporal expression of *Fgf5* strongly supports a stage-specific role for regulating GNP development, particularly along the secondary fissure, to ensure timely differentiation and maturation of granule neurons. Indeed, as we observed in the P0 Sufu-cKO cerebellum, deregulation of *Fgf5* expression correlates with the continued uncontrolled proliferation of GNPs that have failed to differentiate normally. Follow-up studies geared towards characterizing cell-specific expression of *Fgf5* and how FGF5 acts on GNP differentiation are crucial to solidifying the exact function of FGF5 in early neonatal cerebellar development.

Our findings are contrary to previous reports of the proliferation-suppressive roles of FGFs, particularly FGF2, in GNPs specifically carrying *Ptch1* mutations (Fogarty et al., 2007; Emmenegger et al., 2013). However, in contrast to the Ptch1-cKO cerebellum, the newborn Sufu-cKO cerebellum still expresses PTCH1 and exhibits reduced GLI1, GLI2, and GLI3 protein levels (**Supplementary Fig. 1A**). These key molecular differences may activate unique signaling networks in Sufu-cKO GNPs. Additionally, 18 FGF ligands differ significantly in molecular features and binding specificities to distinct combinations of FGFR splice variants (Ornitz and Itoh, 2015). For example, unlike FGF2, which acts in an autocrine manner, FGF5 can exert both autocrine and paracrine functions, bind different combinations of FGFRs, and is dynamically expressed in distinct regions of the developing cerebellum. Thus, in-depth studies are needed to elucidate the exact mechanisms triggered by abnormally high levels of FGF5 in the developing cerebellum, particularly since *Fgf5* overexpression is known to drive cancer development and progression, including brain tumors (Allerstorfer et al., 2008). Importantly, examination of whether inhibition of FGF signaling by AZD4547 (or other FGFR inhibitors) in MB tumor-bearing Sufu;p53-dKO mice is crucial to establish that FGF signaling is a druggable target for SUFU-associated infantile MB^SHH^.

The high occurrence of *SUFU* mutations in infantile MB indicates the selective vulnerability of the developing cerebellum to the neoplastic effects of SUFU dysfunction. Notably, the timing of *SUFU* LOF is critical; conditional deletion of *Sufu* in neural stem cells before granule neuron specification (using the hGFAP-Cre line) results in GNP hyperplasia, whereas conditional deletion of *Sufu* after granule neuron specification (using the Atoh1-Cre line) does not lead to these defects (Jiwani et al., 2020). Thus, *SUFU*-associated infantile MB^SHH^ is a likely consequence of defects stemming from the early stages of granule neuron lineage specification at embryonic stages. However, since hGFAP-Cre may also be expressed in cerebellar glial cells, we cannot yet eliminate the possibility that the defects are directly or indirectly a consequence of abnormal glial cell development and function. In-depth studies are required to interrogate the effects of SUFU LOF in cerebellar glial cell development and how this might indirectly affect GNP differentiation.

Unfortunately, tumors initiated at embryonic stages are typically undetectable until several months after birth, when tumorigenesis has significantly progressed. Thus, therapeutics for infantile MB must successfully curtail tumorigenic mechanisms at postnatal stages and minimally affect normal GNPs elsewhere in the cerebellum. Towards this goal, inhibiting localized FGF5 and FGF signaling activity may provide new paths toward the design of targeted treatments. Since we confirmed the occurrence of high *FGF5* levels in a subset of infantile MB^SHH^ patients, measuring FGF5 may be a useful diagnostic biomarker for this patient population. This may predict the lack of efficacy of SHH-targeting compounds in curtailing tumor growth but could instead significantly impede cerebellar development. Importantly, detection of elevated FGF5 levels may identify patients who will be responsive to FGF-targeting treatments. Ultimately, we hope these studies facilitate the design of much-needed precision medicines to address the distinct oncogenic mechanisms specifically and effectively in infantile MB^SHH^ patients while enabling normal progression of cerebellar development.

## Acknowledgements

We thank Hirofumi Noguchi and other members of the Pleasure Lab for critical discussions, the UCSF Center for Advanced Light Microscopy for assistance with imaging, and William Krause for assistance with transcriptomics. Schematic diagrams were created with BioRender.com. This work was supported by the NIH R01MH077694 and R01NS118995 (S.J.P.), R01MH077694-S1 (H.G.), NIH/NCI K01CA201068 (O.R.Y.), and American Brain Tumor Association Grant #A131363 (O.R.Y.).

This paper was typeset with the bioRxiv word template by @Chrelli: www.github.com/chrelli/bioRxiv-word-template.

## Materials and Methods

### Animals

Mice carrying the floxed *SUFU* allele (Sufu^fl^) were kindly provided by Dr. Chi-Chung Hui (University of Toronto) and were genotyped as described elsewhere (Pospisilik et al., 2010). *hGFAP-cre* (Stock #004600; Schuller et al., 2008) and Gli1-LacZ (Stock #008211) mice were obtained from Jackson Laboratories (Bar Harbor, ME, USA). Mice designated as controls did not carry the *Cre* transgene and may have either one of the following genotypes: *Sufu*^*fl/+*^ or *Sufu*^*fl/fl*^. All mouse lines were maintained in mixed strains, and the analysis included male and female pups from each age group, although sex differences were not included in data reporting. All animal protocols were in accordance with National Institutes of Health regulations and approved by the UCSF Institutional Animal Care and Use Committee (IACUC).

### *In vivo* treatment with FGFR inhibitor

The FGFR1-3 inhibitor, AZD4547 (Selleck Chemicals, #S2801), was dissolved sequentially in 4% Dimethyl Sulfoxide (DMSO), 30% Polyethylene glycol (PEG), 5% Tween-80, and water to make a 1mM solution. 1 μl of 1mM AZD4547, or vehicle only as control, was injected into the lateral ventricle (∼1 mm from the cerebellar midline) of pups for 5 days from P0/P1 using a 2.5 μl syringe (Model 62 RN; Hamilton Scientific). Pups remained with the mother until perfusion at P7 for analysis.

### Immunohistochemistry and LacZ Staining

Perfusion, dissection, immunofluorescence, and LacZ staining were conducted according to standard protocols as previously described (Yabut et al., 2015). Briefly, P0/P1 brain tissues were fixed after dissection by direct immersion in 4% paraformaldehyde (PFA) overnight. P7 and older postnatal brains were fixed by intracardial perfusion with 4% PFA followed by two h post-fixation. All fixed brains were cryoprotected with a 15-30% sucrose gradient overnight before embedding in Optimal Cutting Temperature (OCT) compound for cryosectioning. Cryostat sections were air dried and rinsed 3× in PBS plus 0.2%Triton before blocking for 1 h in 10% normal lamb serum diluted in PBS with 0.2% Triton to prevent nonspecific binding. A heat-induced antigen retrieval protocol was performed on select immunohistochemistry experiments using 10 □ M Citric Acid at pH 6.0. Primary antibodies were diluted in 10% serum in PBS with 0.2% Triton containing 4’6-diamidino-2-phenylindole (DAPI); sections were incubated in primary antibody overnight at room temperature. The following antibodies were used: rabbit anti-Pax6 (1:250 dilution; Cat. #: 901301, Biolegend); rabbit anti-NeuN (1:250 dilution; Cat. #: PA5-784-99, Invitrogen); mouse anti-Calretinin (1:250, Cat. #: AB5054, Millipore); rabbit anti-phospho-Erk1/2 (1:250 dilution; Cat. #: 4370, Cell Signaling); gH2AX (1:100 dilution; Cat. #: 05-636, Millipore); mouse anti-Ki-67 (1:100 dilution; Cat. #: 550809 BD Biosciences) and cleaved-Caspase 3 (1:250 dilution; Cat. #: 9661S, Cell Signaling). To detect primary antibodies, we used species-specific Alexa Fluor-conjugated secondary antibodies (1:500; Invitrogen) in 1X PBS-T for 1 h at room temperature, washed with 1X PBS, and coverslipped with Fluoromount-G (SouthernBiotech).

### In Situ Hybridization

RNAScope ISH was conducted for *Fgf5* and *Ptch1*. RNAscope probes for Mm-*Fgf5* were designed commercially by the manufacturer (catalog # 417091, Advanced Cell Diagnostics, Inc.). RNAScope Assay was performed using the RNAscope Multiplex Fluorescent Reagent Kit V2 according to the manufacturer’s instructions with the following conditions. Slides of cryosectioned brain tissues were prepared by air-drying for 30 minutes at 60ºC, then post-fixed in 4% PFA in 1X PBS for 15 minutes at 4ºC. Tissues were dehydrated at room temperature in an ethanol gradient (50% Ethanol, 70% Ethanol, then 100% ethanol) for 5 minutes each, followed by a final 100% ethanol wash for 5 minutes. RNAScope Hydrogen Peroxide treatment of tissues occurred for 10 minutes. Target retrieval was performed by immersing slides in RNAscope 1X Target Retrieval Reagent for 5 minutes in a 99ºC steamer. Tissue sections were subjected to RNAscope Protease Plus reagent treatment for 10-15 minutes at 40ºC in HybEZ Oven (ACDBio). Tissues were hybridized with the probe mix and incubated for 2 hours at 40ºC in the HybEZ Oven. AMP1, AMP2, and AMP3 sequential hybridization steps were performed as per manufacturer’s instructions. Slides were incubated in RNAScope Multiplex FL v2 HRP-C1 for 15 minutes at 40ºC. Signals were detected using the TSA Plus Fluorescein Kit (catalog # NEL741E001KT, PerkinElmer) and Opal Dye reagent (Opal 570, catalog # FP1488001KT or Opal 520, catalog # FP1487001KT, from Akoya Biosciences) after 30 minutes of incubation at 40ºC.

### Western Blot Analysis

Western blot analyses were conducted according to standard protocols. Soluble extracts were loaded onto Criterion, 4-15% Tris-HCI 4 SDS-PAGE gels (Bio-Rad), separated at 120V, and transferred to PVDF membrane at 30V for 2 hours or overnight at 4°C. Membranes were blocked with 3% milk/1X TBS-T (Tris-buffered saline with 0.1% Tween 20) or 5% BSA/1X TBS-T for 1hr at room temperature and incubated with primary antibodies diluted in blocking buffer overnight at 4°C, and secondary antibodies (1:5000 dilution; IR-Dye antibodies, LI-COR) for 1 hour at RT. Membranes were washed in 1X TBS-T and scanned using the Odyssey Infrared Imaging System (LI-COR). Primary antibodies were used as follows: rabbit anti-Gli1 (1:1000; Abcam); goat anti-Gli2 (1:1000; R&D Systems), rabbit anti-Gli3 (1:100; Santa Cruz), rabbit anti-Pax6 (1:1000 dilution; Cat. #: 901301, Biolegend); rabbit anti-NeuN (1:1000 dilution; Cat. #: PA5-784-99, Invitrogen); rabbit anti-GABA A Receptor α6 (1:1000 dilution; Cat. #. PA5-77403, GABRA6; Invitrogen), and a-Tubulin (1:5000 dilution; Cat. #: ab4074, Abcam). Quantification and analysis were conducted using the Odyssey Image Studio Software (LI-COR). Protein levels were normalized to GAPDH protein levels. Levels of NeuN and GABRA6 were quantified in correlation with Pax6 levels (NeuN/Pax6 or GABRA6/Pax6) to determine the proportion of Pax6+ cells expressing mature granule neuron markers.

### Image Analysis and Acquisition

Images were acquired using a Nikon E600 microscope equipped with a QCapture Pro camera (QImaging), Zeiss Axioscan Z.1 (Zeiss, Thornwood, NY, USA) using the Zen 2 blue edition software (Zeiss, Thornwood, NY, USA), or the Nikon Ti inverted microscope with CSU-W1 large field of view confocal and Andor Zyla 4.2 sCMOS camera. All images were imported in tiff or jpeg format. Brightness, contrast, and background were adjusted equally for the entire image between controls and mutants using the “Brightness/Contrast” and “Levels” functions from the “Image/Adjustment” options in Adobe Photoshop or NIH ImageJ without any further modification. NIH Image J was used to threshold background levels between controls and mutant tissues to quantify fluorescence labeling. To quantify cell density, positively labeled cells within defined EGL regions, as defined in **Fig. 2A**, were counted. Quantification of double-labeled cells was performed in 1-2 mm optical slices obtained by confocal microscopy. We relied on continuous DAPI nuclear staining to distinguish individual cells in each optical slice to determine the cellular colocalization of specific markers being analyzed (e.g., pERK and Ki-67). All measurements were performed from 2-3 20 mm thick and histologically matched cerebellar sections of 3-6 independent mice per genotype analyzed. Individual points in the bar graphs represent the average cell number (quantified from 2-3 sections) from each mouse.

### Human MB Gene Expression

Expression values of FGF5 (ENSG00000138675) were assessed using Geo2R (Barrett et al., 2013) from published human MB subtype expression dataset accession no. GSE85217 (Cavalli et al., 2017). GEO2R is an interactive web tool that compares expression levels of genes of interest (GOI) between sample groups in the GEO series using original submitter-supplied processed data tables. We entered the GOI Ensembl ID and organized data sets according to age and MB subgroup or MB^SHH^ subtype classifications. GEO2R results presented gene expression levels as a table ordered by FDR-adjusted (Benjamini & Hochberg) p-values, with significance level cut-off at 0.05, processed by GEO2R’s built-in limma statistical test. The resulting data were subsequently exported into Prism (GraphPad). Scatter plots presenting FGF5 expression levels across all MB subgroups (Figure 1A) and MB^SHH^ subtypes (Figure 1D). We performed additional statistical analyses to compare FGF5 expression levels between MB subgroups and MB^SHH^ subtypes and graphed these data as violin plots (Figure 1B, 1C, and 1E). For these analyses, we used one-way ANOVA with Holm-Sidak’s multiple comparisons test, single pooled variance. *P* value ≤0.05 was considered statistically significant. Graphs display the mean ± standard error of the mean (SEM). Sample sizes analyzed were: MB^WNT^ n=70, MB^SHH^ n=224, MB^GR3^ n=143, MB^GR4^ n=326, MB^SHHa^ n=66, MB^SHHb^ n=35, MB^SHHg^ n=47, and MB^SHHd^ n=76.

### Statistics

Prism 8.1 (GraphPad) was used for statistical analysis. Two sample experiments were analyzed by Student’s *t* test, and experiments with more than two parameters were analyzed by ANOVA. In 1-or 2-way ANOVA, when interactions were found, follow-up analyses were conducted for the relevant variables using Holm-Sidak’s multiple comparisons test. All experiments were conducted at least in triplicate with sample sizes of n = 3−6 embryos/animals/slices per genotype. *P* value ≤0.05 was considered statistically significant. Graphs display the mean ± standard error of the mean (SEM). Statistical values and analyses are summarized in **Table 1**.

---

**Supplementary Figure 1:**
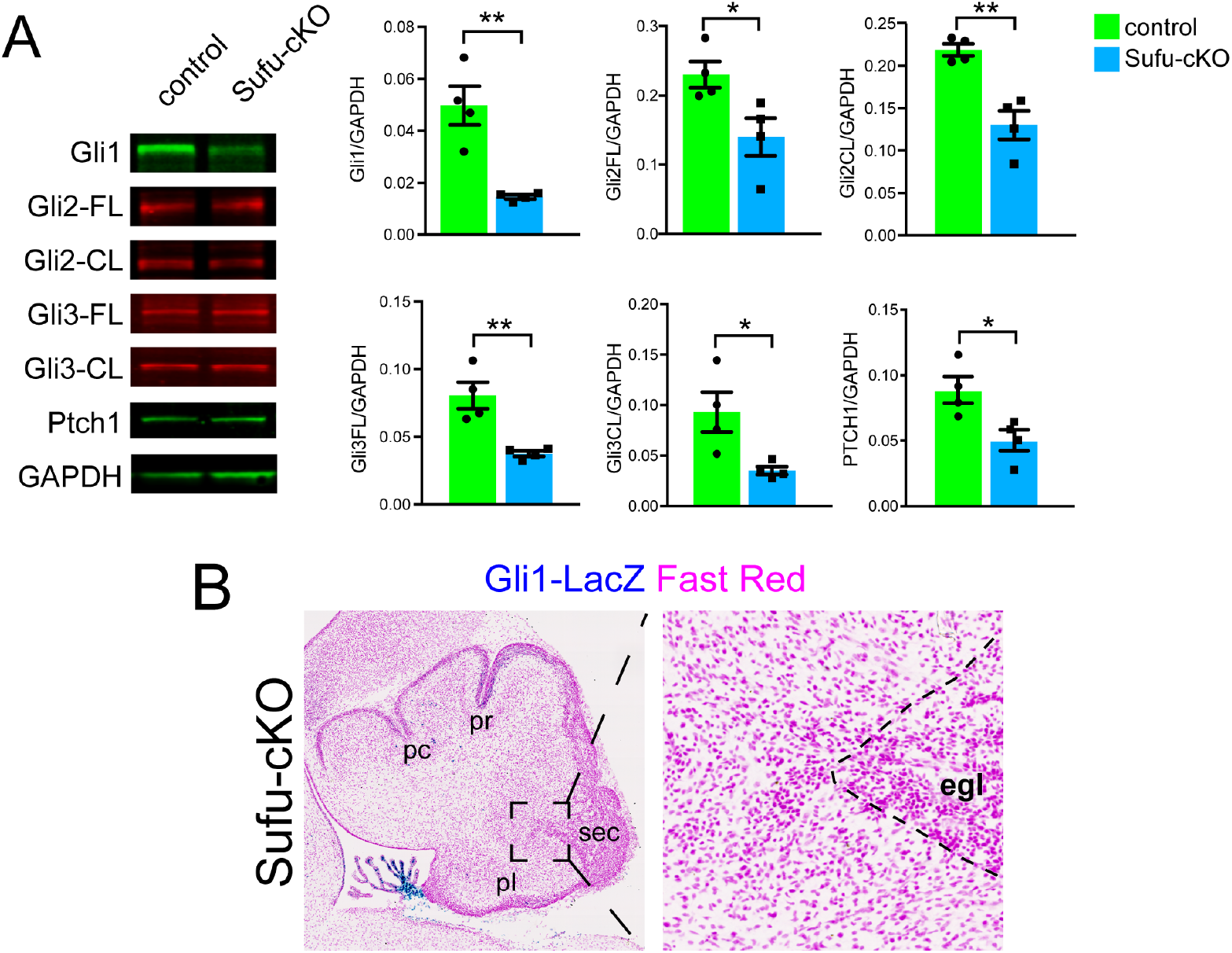
Reduced SHH signaling activity in the P0 Sufu-cKO cerebellum. **(A)** Western blot analysis of cerebellar protein lysates from P0 control and Sufu-cKO mice showing significantly lower levels of total and cleaved versions of Gli transcription factors in the P0 Sufu-cKO cerebellum. *p<0.05, **p<0.01 (**B)** b-galactosidase activity (blue), representing the Gli1-LacZ transgene, is largely absent in areas adjacent to the EGL along the secondary (sec) fissure of the P0 Sufu-cKO cerebellum.

**Supplementary Figure 2:**
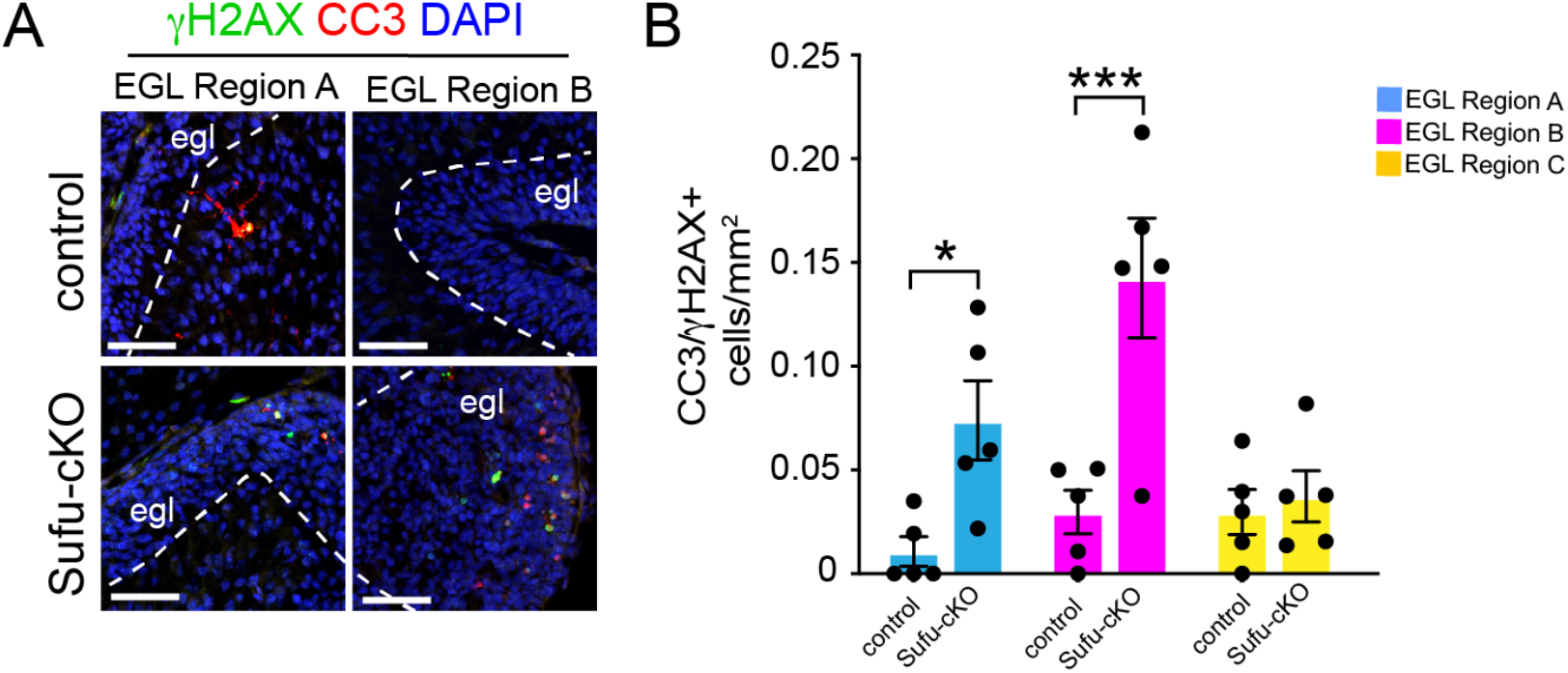
Evidence of pre-neoplastic lesions and high rates of cell death in Sufu-cKO granule neuron precursors. **(A)** Double-immunofluorescence staining with (γH2AX (green) and cleaved-caspase 3 (CC3; red), a marker for apoptotic cells, and DAPI labeling in regions A and B. Scale bars = 50 μm **(B)** Quantification of the density of cells labeled with CC3 and (γH2AX within each EGL regions. \**p*<0.05, ***p<0.001.

